# Evaluation of ERIC-PCR and MALDI-TOF as typing tools for multidrug resistant *Klebsiella pneumoniae* clinical isolates from a tertiary care center in India

**DOI:** 10.1101/2022.07.06.499023

**Authors:** Jyoti Kundu, Shubhangi Kansal, Shivali Rathore, Meenakshi Kaundal, Archana Angrup, Manisha Biswal, Kamini Walia, Pallab Ray

## Abstract

**Background and Aim:** Multidrug resistant *Klebsiella pneumoniae* is associated with nosocomial infections in both outbreak and non-outbreak situations. The study intends to evaluate the potential of enterobacterial repetitive intergenic consensus-polymerase chain reaction (ERIC-PCR), a genomic based typing and matrix-assisted laser desorption ionization time-of-flight mass spectrometry (MALDI-TOF MS) proteomic-based typing techniques for clonal relatedness among multidrug resistant *Klebsiella pneumoniae* isolates.

**Methodology:** Multidrug resistant clinical isolates of *Klebsiella pneumoniae* (*n* =137) were collected from March 2019 to February 2020. Identification and protein-based phylogenetic analysis were performed by MALDI-TOF MS. Genomic typing was done by ERIC-PCR and analyzed by an online data analysis service (PyElph). Dice method with unweighted pair group method with arithmetic mean (UPGMA) program was used to compare the ERIC profiles. The samples were also evaluated by PCR for the presence of genes encoding carbapenemases, extended spectrum beta lactamases (ESBLs) and mobile colistin resistance-1 (*mcr1*).

**Result and Conclusion:** Isolates were typed into 40 ERIC types, and six groups by MALDI-TOF-MS. PCR-based analysis revealed that all the strains harbored two or more ESBL and carbapenemase genes. None of the isolates revealed the presence of the plasmid mediated *mcr-1* gene for colistin resistance. The study presents ERIC based typing as more robust in comparison to MALDI-TOF for finding the clonal relatedness in epidemiological studies.

## Introduction

*Klebsiella pneumoniae* is an opportunistic gram-negative bacterium accounting for approximately one third of all hospital and community acquired infections. These include pneumonia, surgical wound infections, meningitis, urinary tract infections and blood stream infections (1). Increase in the incidence and prevalence of multidrug resistant (MDR) *K. pneumoniae* has been associated with therapeutic failures, prolonged hospitalization, high mortality rates and a significant economic burden (2). In the last two decades, there has been a tremendous increase in infections caused by multiple drug resistant gram-negative bacteria, including those resistant to the last resort drugs like carbapenems and colistin (3,4). The World Health Organization has prioritized *Klebsiella pneumoniae* as a target for the development of newer antimicrobials for the treatment of nosocomial infections (5). Several transmission dynamics studies in both outbreak and non-outbreak situations for ESBL-producing *K. pneumoniae* infections have shown the presence of high clonal diversity (6–8).

Molecular typing techniques are powerful tools used to determine clustering among multidrug resistant (MDR) *K. pneumonia*e and to derive information about their transmission. We used enterobacterial repetitive intergenic consensus-polymerase chain reaction (ERIC-PCR) and matrix-assisted laser desorption ionization time-of-flight (MALDI-TOF) to assess clonality of *K. pneumoniae* isolates. MALDI typing relies on the microbial proteomic spectra (fingerprints) clustering, whereas ERIC-PCR relies on the fingerprints due to the presence of multiple copies of conserved consensus sequences in the genomes of bacteria for epidemiological surveillance (9).

## Material and Methods

### Bacterial Isolation and Identification

A total of 137 non-duplicate MDR *K. pneumoniae* isolates were collected from blood, respiratory, and abscess specimens received in the Department of Medical Microbiology, Postgraduate Institute of Medical Education and Research, Chandigarh, India (PGIMER) from March 2019 to February 2020. Isolates were confirmed as *K. pneumoniae* by MALDI -TOF MS (Bruker Daltoniks, Bremen GmBH Germany). Samples were cultured and incubated overnight at 37°C. Information regarding the type of sample (bronchoalveolar lavage, endotracheal aspirate, sputum, blood, wound swab, and abscess drainage), age and sex of patients and type of admission (inpatient/ward/unit/outpatient) were extracted from the hospital information system. Institutional ethical approval to carry out the study has been obtained (IEC-11/2018-1048).

### Antimicrobial susceptibility testing

Antimicrobial susceptibility of the isolates was performed by the VITEK 2 system against ceftazidime, ceftriaxone, cefepime, ertapenem, imipenem, meropenem, aztreonam, ampicillin-sulbactam, piperacillin-tazobactam, ciprofloxacin, levofloxacin, amikacin, gentamicin, tigecycline and trimethoprim-sulfamethoxazole. The broth microdilution method was performed for minimum inhibitory concentration (MIC) of colistin. *Escherichia coli* ATCC 25922 was used as quality control strain for antibiotic susceptibility tests. Results were interpreted following Clinical and Laboratory Standards Institute (CLSI) guidelines (M-100, Ed 2019) (10). Strains not susceptible to at least one agent in three or more antimicrobial classes were defined as multidrug-resistant and were used in the study.

### Detection of genes conferring resistance to carbapenems, β-lactams and mobile colistin resistance-1

Total DNA of all the isolates of *K. pneumoniae* was extracted using the boiling method and further used to detect the presence of the following genes – *bla*_NDM_, *bla*_KPC_, *bla*_OXA1_ and *bla*_OXA48_, *bla*_VIM,_ *bla*_CTXM-15_, *bla*_SHV_, *bla*_TEM_, and *bla*_IMP_. PCR assays were carried out using specific primers as described by Dallenne *et al* (11) and detection of plasmid mediated colistin resistance was performed using universal primer for CLR5 region of *mcr*1 gene. *K. pneumoniae* ATCC 1705 was used as a standard positive control strain for *bla*_SHV_ and *bla*_KPC_. For the other genes, in-house strains known to harbor the genes (and confirmed by targeted Sanger sequencing) were used as positive controls. A non-ESBL producing organism (*E. coli* ATCC 25922) was used as a negative control. Primers used for amplification are listed in supplementary file.

### Enterobacterial Repetitive Intergenic Consensus-Polymerase Chain Reaction (ERIC-PCR)

ERIC-PCR technique was carried out in thermocycler (Thermo ABI Veriti™ Biosystems) using ERIC primers-forward: 5’-ATG TAA GCT CCT GGG GAT TCAC-3’ and reverse: 5’-AAG TAA GTG ACT GGG GTG AGC G3’. The PCR protocol consisted of an initial denaturation cycle at 95°C for 5 min, followed by 30 cycles of denaturation at 94°C for 1 min, annealing at 48°C for 1 min, extension at 72 °C for 1 min, and a final cycle of amplification at 72°C for 10 min. PCR products were loaded on 1.5% agarose gel (Lonza SeaKem LE Agarose) at constant voltage of 70V for one hour, and the banding patterns were visualized under ultraviolet radiation (supplementary file). ERIC patterns were analyzed by online data analysis service of PyElph (12). ERIC profiles were compared using Dice similarity matrix coefficient and clustered by neighbor joining method to prepare the phylogenetic tree. Isolates with two or more different bands in ERIC banding pattern were considered as different ERIC type. The dendrogram was drawn according to the clusters (Fig1).

**Fig 1:**
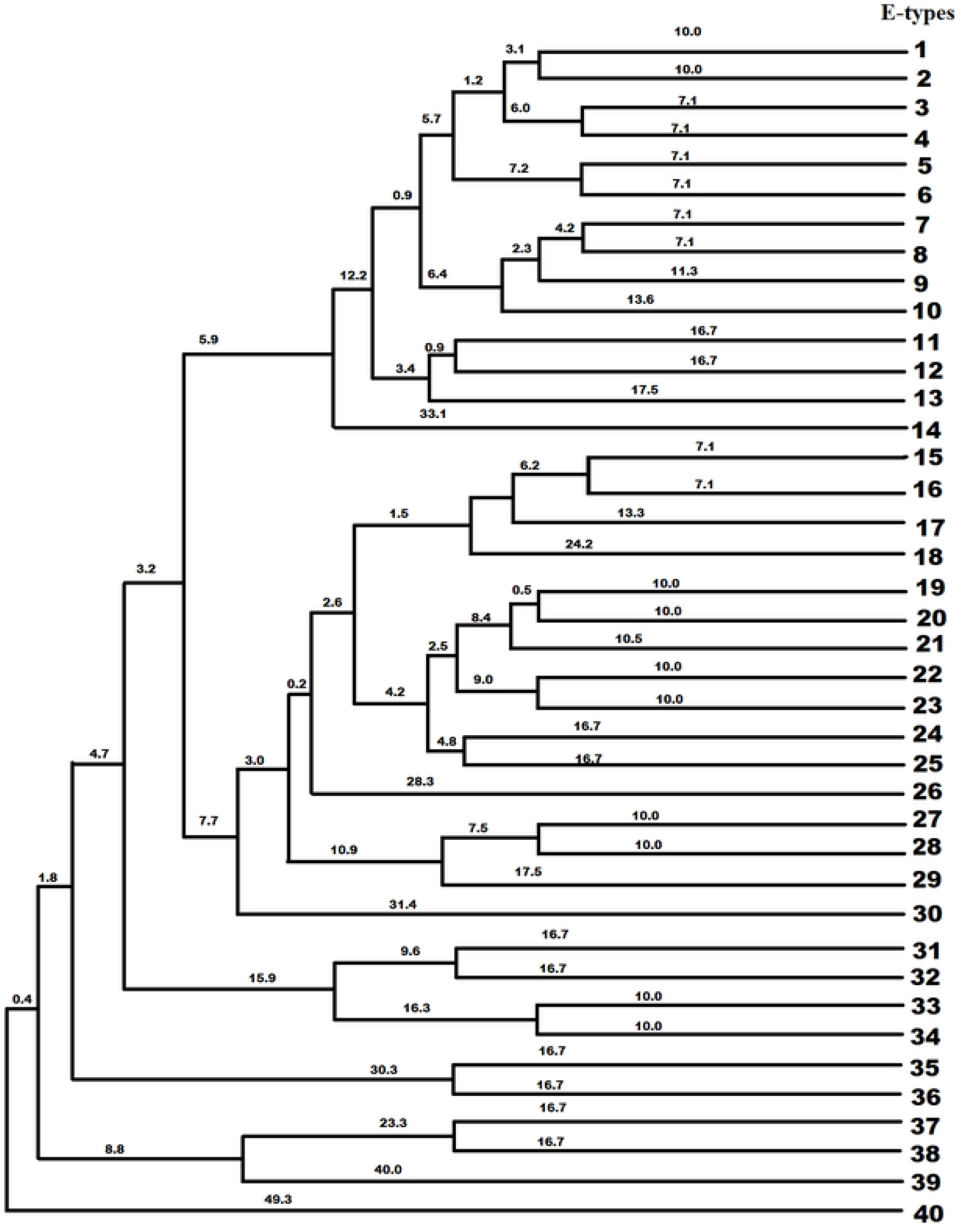
Dendrogram generated with neighbor end joining and the UPGMA clustering methods, showing the genetic similarity among *K. pneumoniae* isolates by enterobacterial repetitive intergenic consensus (ERIC) genotyping.

### Matrix-assisted laser desorption ionization time-of-flight (MALDI-TOF) typing

Identification and phylogenetic analysis of all the isolates was performed according to the method prescribed by Bruker Daltonik, GmbH. In brief, a single bacterial colony was homogenously smeared onto a target spot on a reusable steel plate (MSP 96 target polished steel BC) using a sterile wooden toothpick. Bacterial components were crystalized by overlaying each spot with 1μl of saturated α-cyano-4-hydroxycinnamic acid matrix solution (10mg/ml conc. of HCCA powder in 50% acetonitrile with 2.5% trifluoroacetic acid). Each spot was air-dried at room temperature. Mass spectra of all the isolates were acquired automatically using default identification settings of FlexControl Software with linear positive mode, mass range 2-20kDa. The final spectrum was the sum of 20 single spectra, each obtained by 200 laser shots on random target spot positions. The raw spectrum obtained were processed and classified with the help of MALDI BioTyper software, version 3.1. The standard interpretative criteria for Bruker were applied. A phylogenetic tree (Fig 2) was constructed from main spectrum profiles of all the isolates using FlexControl software (version 3.4.127.0) and BioTyper software (version 3.1).

**Fig 2:**
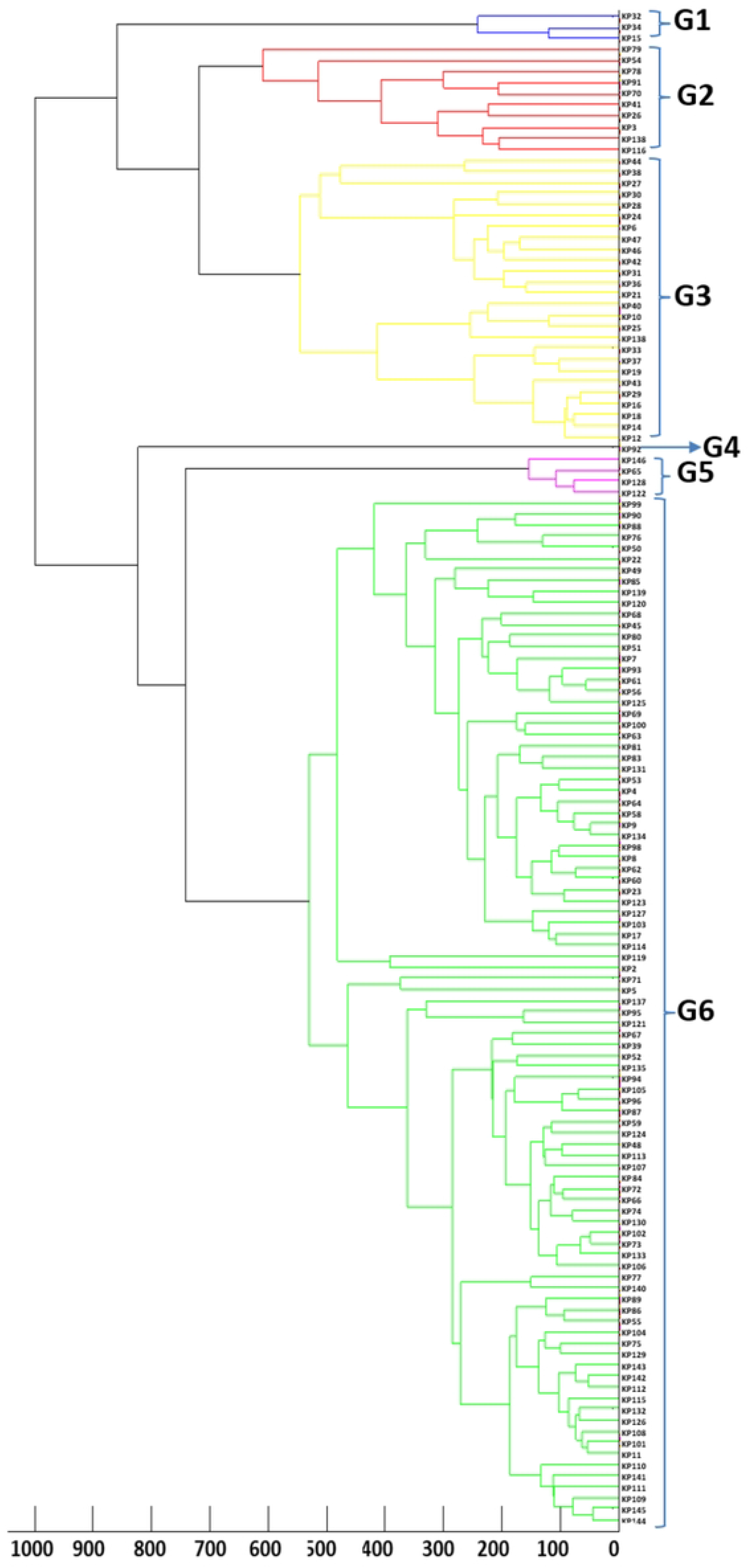
**MALDI-TOF dendrogram-** phylogenetic tree was constructed from main spectrum profiles of all the isolates using Flexcontrol software (version 3.4.127.0) and Biotyper software (version 3.1). Colour coding indicates the MALDI types of the isolates.

### Discriminatory index (D)

The discriminatory indices (D) of MALDI-TOF and ERIC-PCR were calculated based on Simpson’s Index of Diversity formula (13) as below:

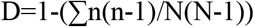

Where D is diversity index, N is the total number of individual strains in the sample population and n is the number of isolates of a particular type/cluster. Simpson’s index of diversity ranges from 0.0 to 1.0 where value equal to 1.0 is highly discriminatory, and a value of 0.0 represents that all the isolates are of an identical type.

## Results

### Clinical isolates

Total 137 multidrug resistant *K. pneumoniae* clinical isolates from endotracheal aspirates, sputum and bronchoalveolar lavage (n=30), wound pus (n=62), body fluids, CSF (n= 7), and blood (n=38) were included (Fig 3). Lactose fermenting colonies on MacConkey agar were further identified by MALDI-TOF MS.

**Fig 3:**
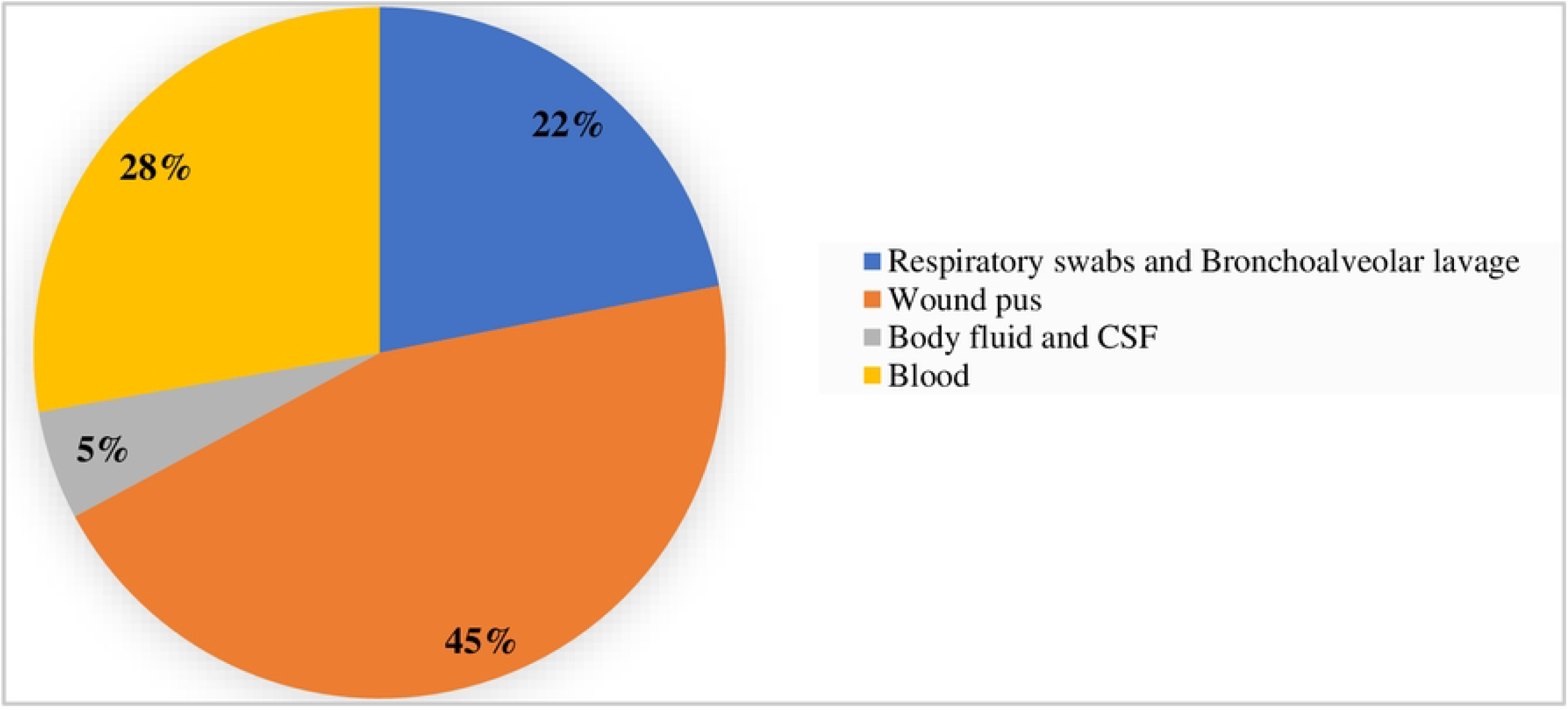
Distribution of clinical isolates as per the origin of sample collection.

### Polymerase chain reaction-based detection of extended spectrum beta-lactamases and carbapenemase-encoding genes

Among the various genes tested, all the isolates expressed at least two antimicrobial resistance genes, 95% isolates expressed *bla*_TEM_ followed by *bla*_CTXM-15_ (∼88%). Amongst the carbapenemase genes tested, maximum expression was for *bla*_OXA_ (∼88%) followed by *bla*_NDM_ (∼54%) as indicated in Table1 and Fig 4.

**Table 1:**
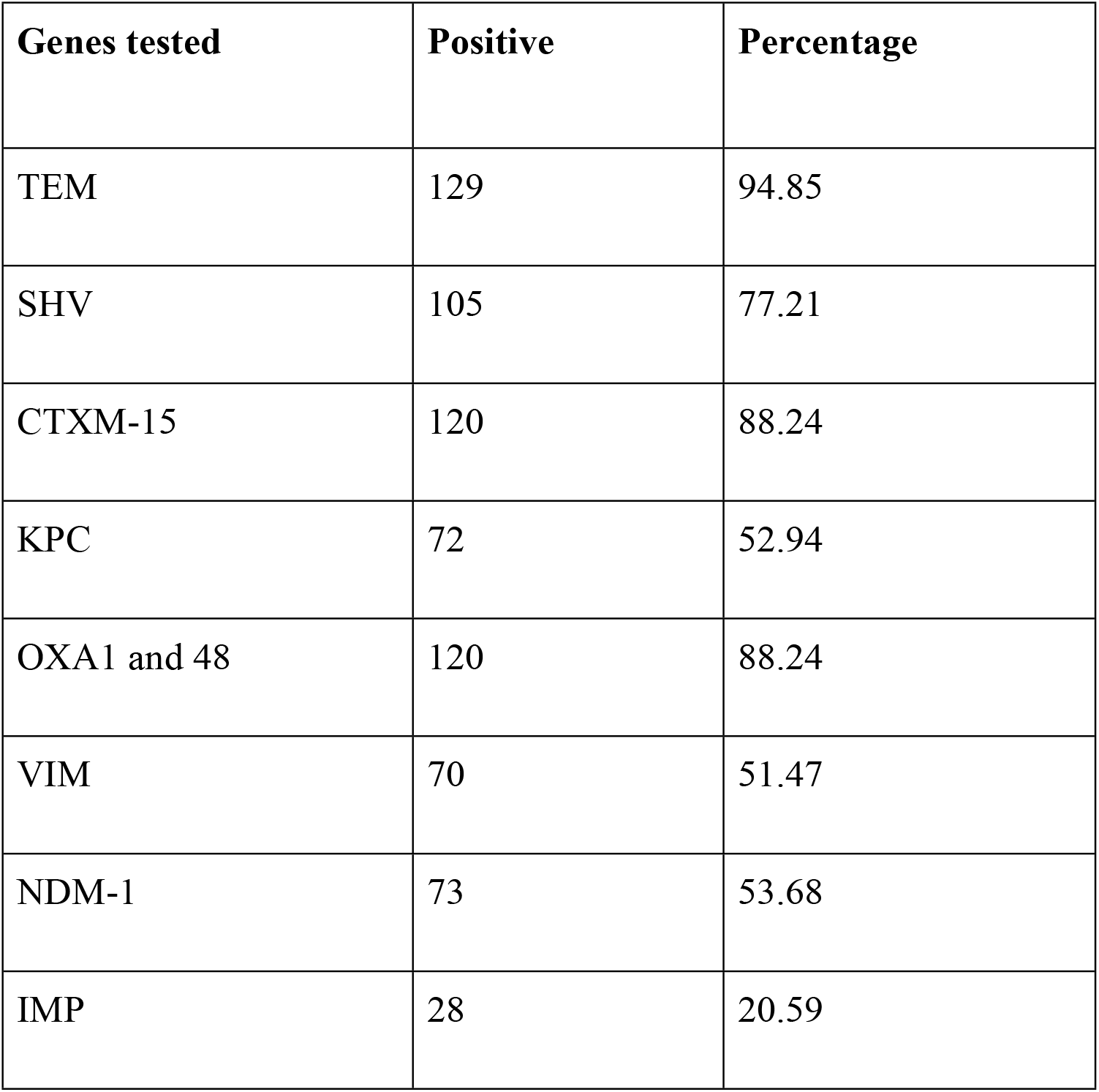
PCR based ESBL and carbapenemase gene expression analysis.

| Genes tested | Positive | Percentage |
| --- | --- | --- |
| TEM | 129 | 94.85 |
| SHV | 105 | 77.21 |
| CTXM-15 | 120 | 88.24 |
| KPC | 72 | 52.94 |
| OXA1 and 48 | 120 | 88.24 |
| VIM | 70 | 51.47 |
| NDM-1 | 73 | 53.68 |
| IMP | 28 | 20.59 |

**Fig 4:**
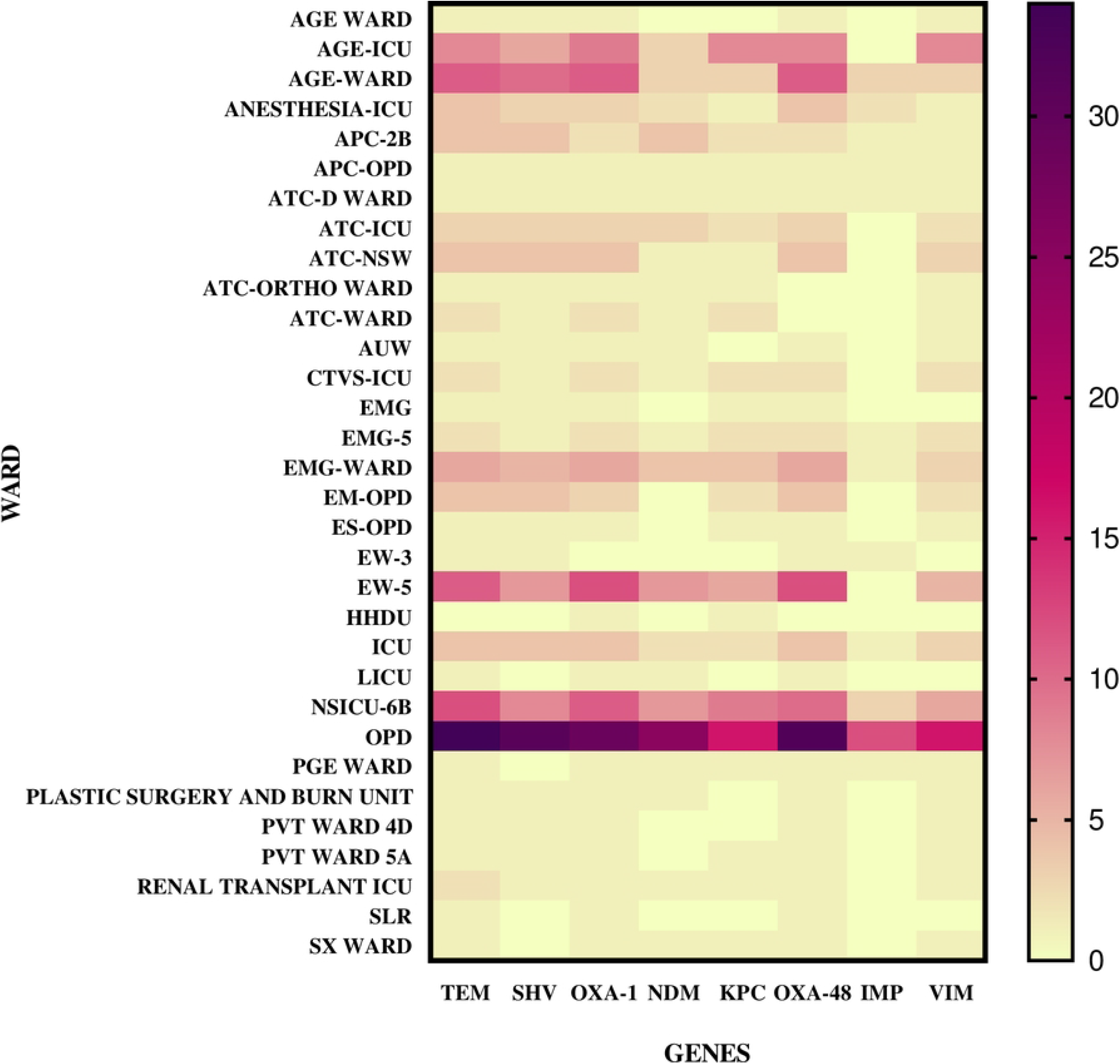
Heat map representing the carbapenemases/ESBL gene distribution with respect to isolates from different hospital units/wards.

### Molecular fingerprinting by ERIC-PCR

The number of bands varied from 1 to 11 with the size ranging from 100 bp to more than 1.5kb. A total of 40 different ERIC profiles (E-types) were observed designated as E1 to E40. Out of 137 isolates, total 44 isolates belonged to E1, 8 isolates to E2,17 isolates to E11, 12 isolates to E19 type, 2 each to E5, E6, E12, E13, E16, E31, E33, E24, E10=6 isolates, E7 and E35=4 each, E21=3 and other remaining 24 isolates showed a unique pattern. To further investigate probable outbreak of MDR *K. pneumoniae* infection, ERIC typing data was mapped in relation to hospital unit/ward (Fig 5). Details of the same has been provided in the supplementary file.

**Fig 5:**
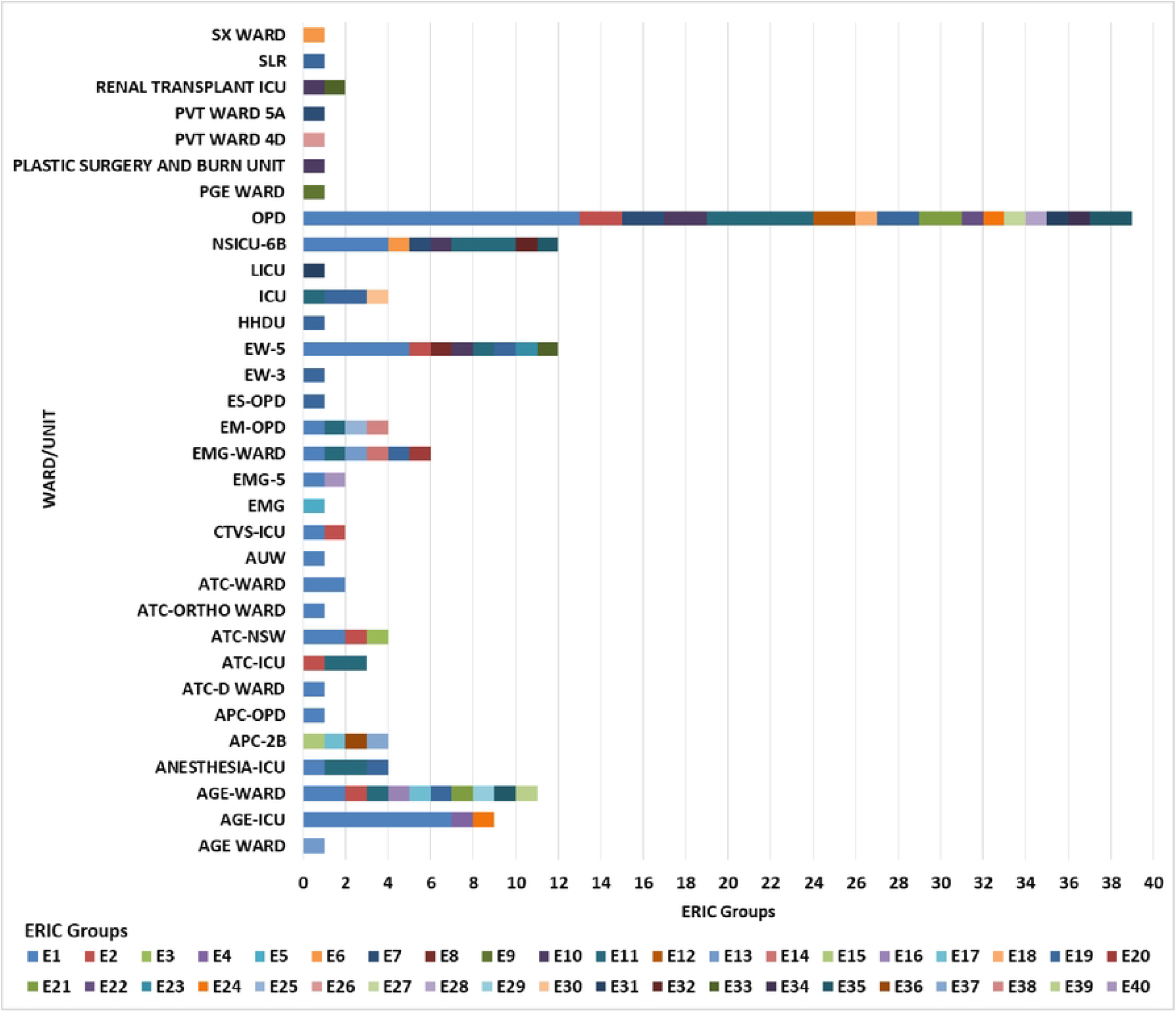
ERIC typing of the isolates as per the sample collection site/ward/unit.

### Matrix assisted laser desorption ionization time of flight analysis

Isolates were allotted into 6 groups *i*.*e*., G1to G6 by MALDI-TOF analysis. G1=3 isolates, G2=10 isolates, G3=26 isolates, G4=1 isolate, G5=4 isolates and G6=93 isolates.

### Discriminatory potential of ERIC-PCR and MALDI-TOF in typing of *K. pneumoniae* isolates

The Simpson’s Diversity Index was calculated for ERIC-PCR, and MALDI-TOF analysis as depicted in the table below. ERIC -PCR was found to be more discriminatory (D=0.8704) for clonal relatedness and was a better phylotyping tool in comparison to MALDI-TOF typing (D=0.5001).

| Typing method | Simpson's Discriminatory index |
| --- | --- |
| ERIC-PCR | 0.8704 |
| MALDI-TOF | 0.5001 |

## Discussion

Given the high prevalence of MDR *K. pneumoniae*, it is essential to understand and establish the clonal relatedness among isolates to prevent and control *K. pneumoniae* outbreaks in a healthcare setting. Various molecular typing methods such as multilocus sequence typing (MLST), pulsed-field gel electrophoresis (PFGE), plasmid typing, antibiogram typing and ribotyping have been used for epidemiological and surveillance studies. However, many of these are expensive, labor intensive and time-consuming. Methods such as palindromic repetitive element-based ERIC-PCR and proteomic signature based MALDI-TOF are quick, reliable, and cost-effective techniques for molecular typing of the Enterobacteriaceae family. We therefore studied these tools to determine the transmission dynamics of MDR *K. pneumoniae* in hospital settings.

*Klebsiella pneumoniae* is a problem pathogen with a high incidence of multidrug resistant strains. To study the epidemiological relatedness among the MDR *K. pneumoniae* isolates and to find out the possibility of any plausible outbreak, we designed this study comparing two methods for typing, one based on the genotype (ERIC-PCR) and another on the basis of protein profiling (MALDI-TOF). The importance of molecular typing of MDR-*K. pneumoniae* by these methods is in strengthening the epidemiological surveillance and to recognize the clonal spread in a rapid and cost-effective manner in comparison to WGS based surveillance. We analyzed a total of 137 isolates and found the genotyping-based method to be more robust and discriminatory than MALDI-TOF as indicated by the Simpson’s diversity index value (for ERIC-PCR, D=0.8704).

In this study, 45% isolates were related to wound pus followed by 28% from blood and 22% from respiratory specimens (Fig 3). ERIC-PCR devised phylogenetic group E1 was most prevalent (29%) and represented a dominant clone, followed by E11 (12.4%) and E19 (8.7%). Out of the 40 isolates from E1 phylogenetic group 13 isolates were collected from general outpatient department (OPD), 7 from AGE-ICU and rest of the isolates from other indoor and outpatient hospital units/wards. Similarly, isolates belonging to other ERIC phylogenetic groups were sampled from different indoor and outpatient hospital units/wards. Our observations revealed that no outbreak or nosocomial clustering happened during the designated period of sampling.

MALDI typing showed a group G6 (68%) was most prevalent followed by G3 (19%). The unit/ward data also did not show any correlation with MALDI groups; supporting our observation from ERIC-PCR data. Both the techniques indicated the convergent result of no likely outbreak/nosocomial spread of MDR *K. pneumoniae* isolates during the study period. Although both the tools provided convergent outcomes (Fig 6), ERIC-PCR was a better tool with Simpson’s diversity Index (D=0.8704) approaching close to numerical value of one. Of the 137 MDR *K. pneumoniae* isolates used in this study, ERIC-PCR revealed 40 (E1-E40) and MALDI-TOF revealed 6 (G1-G6) distinct groups. Among the isolates that were classified as very closely related based on their MALDI-TOF dendrogram, the ERIC-PCR banding patterns showed lack of genetic relatedness, thereby highlighting the difference in clustering the isolates based on genomic versus proteomic signatures. The large number of serotypes in this species could also explain this genetic diversity highlighted by the ERIC-PCR genotypic analysis. Similar to other studies, in our study the clustering pattern by MALDI-TOF was different compared to that seen with ERIC-PCR based techniques (14,15). This might be because of the fact that phenotype expressed is not always the true representative of genotype.

**Fig 6:**
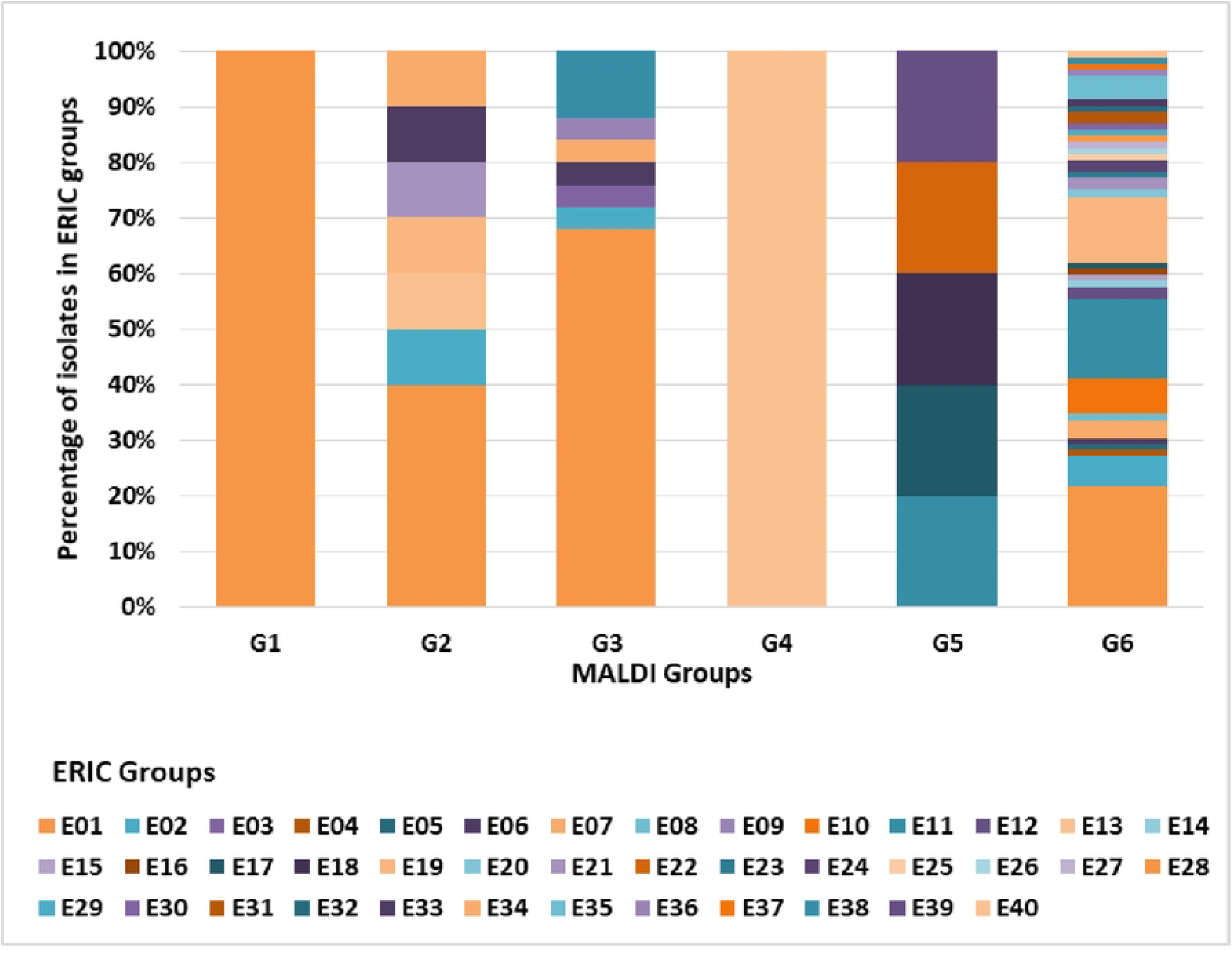
Convergent results with ERIC and MALDI-TOF as typing tools for relatedness of clinical isolates. As depicted in the bar graph ERIC has better discriminatory power for large sample size in comparison to MALDI-TOF.

In this study, the discriminatory indices for ERIC-PCR and MALDI-TOF were performed. Comparison between these two methods revealed the higher discrimination index for ERIC-PCR than MALDI -TOF, but results were not comparable. ERIC-PCR banding pattern gave a meaningful clustering with a Simpson’s Discriminatory index of 0.8704. Proteomics based MALDI-TOF clustering showed that the *Klebsiella pneumoniae* isolates tested were heterogeneous and clonal relatedness could not be depicted as Simpson’s Discriminatory index was 0.5001. Similar to our study Rim et al found MALDI-TOF to have low/insufficient discriminatory power to determine the relatedness of MDR-*Acinetobacter baumannii* isolates (16). A recent study by Purighalla S et al., have the supporting results indicating ERIC being more reproducible and better tool than MALDI-TOF to determine the relatedness of nosocomial *K. pneumoniae* isolates (14) though with routine use of MALDI for microbial identification, it is also being used for outbreak investigations with no added cost for effective interventions (17,18). Although MALDI-TOF is a quick and easy technique and can speed up the infection control measures to prevent further outbreak (19), it does not fit in every situation for perfect discrimination (20). Maskit Bar-Meir also supported the integration of MALDI-TOF-MS, primarily for species identification and only secondarily for epidemiological typing (21). Various studies show that MALDI biotyper has limited ability to provide protein fingerprinting because it targets ribosomal proteins in the limited range of 2-20 kD which is adequate for identification but shows restricted ability to differentiate isolates to the level of clonal complex. Also, there are scopes of improvement for MALDI-TOF MS as a tool for biotyping as its performance also varies based on modification of the growth conditions and extraction method (21, 22).

## Conclusion

Although genomic tools have good discriminatory power but are costly, labour intensive and demand expert handling. ERIC -PCR profiles provide good discriminatory power to find out the clonal relatedness to describe any likely epidemiological outbreak for MDR *K. pneumoniae* isolates. This phylotyping tool is easy, fast, cost effective and can be easily adapted to high throughput analysis of isolates for meaningful clustering.

## Funding disclosure and Acknowledgements

This research work was partially funded by ICMR, India, New Delhi, Ref. No. AMR/158/2018-ECD-II /2020.

## Author’s role

JK: Conceptualization, data curation/analysis and visualization, Writing -Original Draft Preparation, Conducting experimentation, Validation.

SK: Data curation/analysis and visualization, manuscript writing-reviewing/editing, validation. MK: Sample collection and processing.

SR: Sample collection, conducting experimentation, reviewing.

AA: Project administration, Reviewing and editing the manuscript, Data validation, Supervision. MB: Reviewing and editing the manuscript, supervision.

KW: Reviewing the manuscript, Data validation, supervision

PR: Supervision, Project administration, Reviewing and editing the manuscript, Resources.

